# CDK8 and CDK19 kinases have non-redundant oncogenic functions in hepatocellular carcinoma

**DOI:** 10.1101/789586

**Authors:** Katarina Bacevic, Susana Prieto, Stefano Caruso, Alain Camasses, Geronimo Dubra, José Ursic-Bedoya, Anthony Lozano, Jacqueline Butterworth, Jessica Zucman-Rossi, Urszula Hibner, Daniel Fisher, Damien Gregoire

## Abstract

Hepatocellular carcinoma (HCC) is a common cancer with high mortality. The limited therapeutic options for advanced disease include treatment with Sorafenib, a multi-kinase inhibitor whose targets include the Mediator kinase CDK8. Since CDK8 has reported oncogenic activity in Wnt-dependent colorectal cancer, we investigated whether it is also involved in HCC. We find that CDK8 and its paralogue CDK19 are significantly overexpressed in HCC patients, where high levels correlate with poor prognosis. Liver-specific genetic deletion of CDK8 in mice is well supported and protects against chemical carcinogenesis. Deletion of either CDK8 or CDK19 in hepatic precursors had little effect on gene expression in exponential cell growth but prevented oncogene-induced transformation. This phenotype was reversed by concomitant deletion of TP53. These data support important and non-redundant roles for mediator kinases in liver carcinogenesis, where they genetically interact with the TP53 tumor suppressor.

## Introduction

The cyclin-dependent kinase CDK8 is the principal catalytic subunit of the Mediator complex kinase module (1). In mice, genetic deletion of CDK8 or its activating subunit, cyclin C, is embryonic lethal (2, 3). Vertebrate genomes encode a second paralogue of CDK8, CDK19, which also binds cyclin C and can replace CDK8 in the kinase module of Mediator (4, 5). CDK8 may act as an oncogene in several tumor types, including melanoma and colorectal cancer (CRC) (6, 7). Amplification of the CDK8 gene was found in 14% of CRC patients, and reversing CDK8 over-expression in CRC cells with amplified CDK8 reduced cell proliferation by interfering with the β-catenin pathway (6, 8). CDK8 was also uncovered in a mouse transposon-mediated mutagenesis screen for genes whose alteration contributes to intestinal cancer (9). However, intestine-specific *Cdk8* deletion in mice failed to confirm an oncogenic role in intestinal tumorigenesis triggered by mutation of the tumor suppressor *Apc* ((10), and our unpublished results).

CDK8 acts as a co-factor of many transcriptional activators and participates in regulating the expression of a large number of genes, including immediate early genes (11–14), targets of β-catenin (6), p53 (15), c-Myc (16), Hypoxia-inducible factor 1α (HIF1α) (17, 18), Nuclear-factor kappa B (NFκB) (19) and Notch1 (3, 20). CDK8 expression appears to maintain tumors in an undifferentiated state by regulating c-Myc programmes (16). CDK8 was also identified as a crucial regulator of tumor-promoting activity of senescent cells (21). It is presently unclear how these multiple pathways participate in tumor promotion by CDK8, and whether its paralogue CDK19 has similar roles in tumorigenesis.

Since the oncogenic roles of CDK8 in Wnt-dependent CRC have been well documented in the context of constitutive Wnt signalling, we wondered whether it also has roles in hepatocellular carcinoma (HCC), another tumor type frequently associated with activation of Wnt/β-catenin signalling (22). HCC is the main primary liver tumor and one of the most deadly cancers worldwide (23). Mutations in the *CTNNB1* gene, coding for β-catenin, are found in around 30% of patients and are mainly associated with the well differentiated, less aggressive, class of HCC (groups G5 - G6) (22, 24). CDK8 is expressed in the liver, where it may regulate lipogenesis (25). Importantly, lipid accumulation as well as *de novo* lipid biosynthesis and the resulting lipotoxicity lead to hepatic inflammation, constituting major risk factors for liver tumorigenesis (22).

In this study, we show that although CDK8 has a limited role in liver homeostasis both CDK8 and CDK19 paralogues are required for hepatic carcinogenesis in the context of wild-type p53.

## Results

### *CDK8* and *CDK19* are highly expressed in p53-mutated hepatocellular carcinoma

To test whether expression of CDK8 or that of its paralogue CDK19 is altered in hepatic carcinogenesis, we first quantified by qRT-PCR their expression in a large cohort of HCC patients (n=268, Supplemental Table 1). We found that both CDK8 and CDK19 are significantly overexpressed in HCC tumors compared to non-tumoral counterparts or normal liver (Figure 1A). Moreover, we detected a correlation between the expression of the two kinases in HCC (Figure 1B), as previously observed in breast cancer (26). Analysis revealed no correlation with a specific aetiology (alcohol, viral infection, metabolic syndrome). However, there was a highly significant difference in CDK8/19 expression among the HCC subgroups defined by the classification based on clinical and molecular features (24, 27): CDK8 or CDK19 high expressors were enriched in the aggressive G1-G3 subsets as compared to G4-G6 (Figure 1C). Coherently, high CDK8 or CDK19 expression was correlated with mutant p53 status, molecular prognostic 5-gene score (28) and macroscopic vascular invasion (Figure 1D, E and F). Finally, high level expression of CDK8 or CDK19 correlated with poor prognosis (Figure 1G). Thus our data are consistent with an oncogenic role for Mediator kinases in hepatocellular carcinoma.

**Figure 1:**
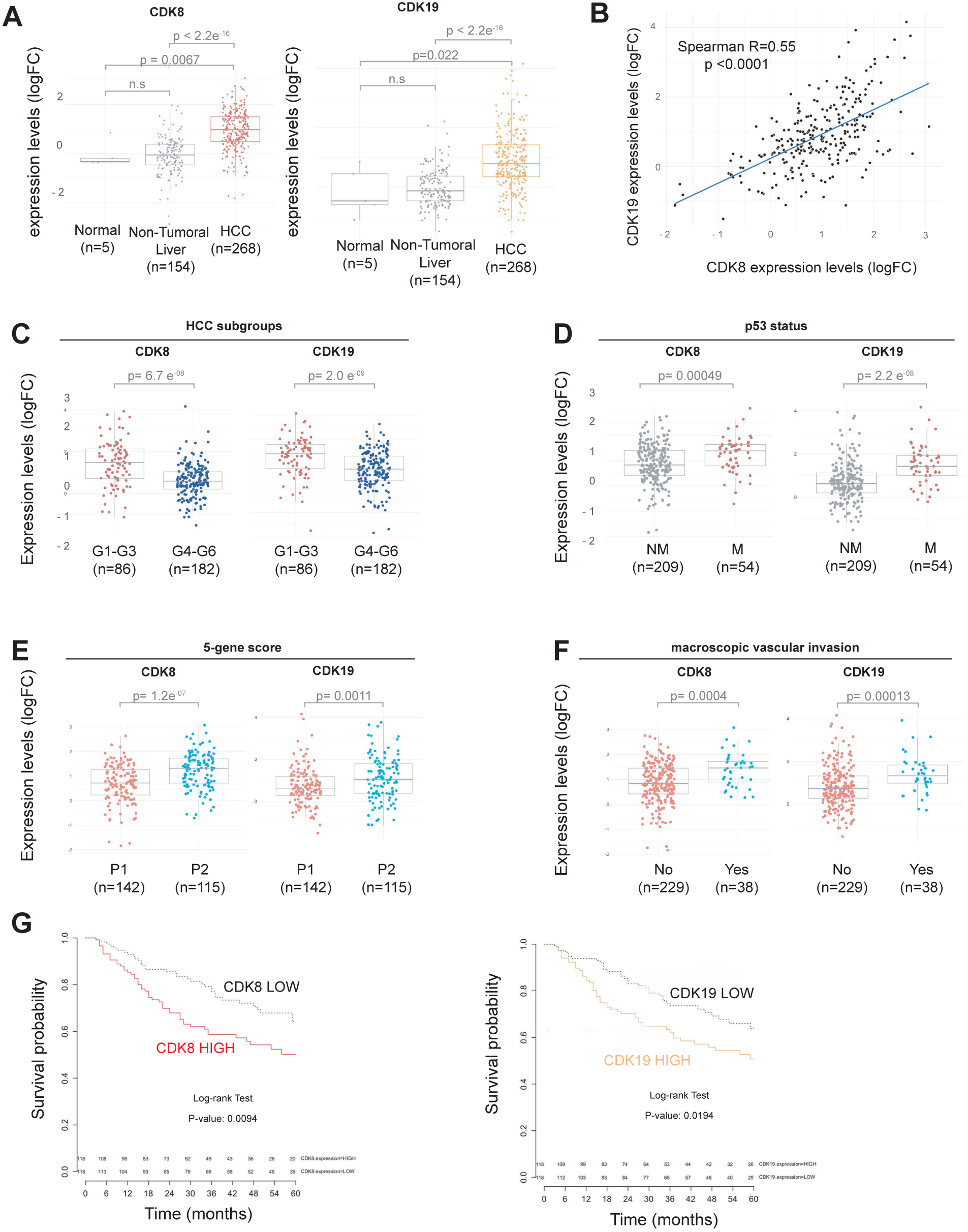
CDK8 and CDK19 are highly expressed in human HCC and associated with mutated p53 status. (**A**) Levels of CDK8 and CDK19 mRNA quantified by qPCR in control liver, non-tumoral part or malignant (HCC) tumor of patients. The fold change (LogFC) in gene expression is presented relative to mean expression level of the corresponding gene in five normal liver samples. Data are represented as Tukey’s boxplots where box indicates the first and third quartiles, bar indicates median, whiskers indicate 1.5 interquartile range (IQR). P-values obtained with the Wilcoxon rank-sum test are indicated. (**B**) Correlation between the expression of CDK8 and CDK19 mRNA levels in human HCC samples (n=268) using Spearman’s rank-order correlation (**C-F**) Levels of CDK8 and CDK19 mRNA in HCC classified according to molecular classification of HCC (Boyault et al., 2007), mutated status of p53, 5 gene score (Nault et al., 2013), or presence of macroscopic vascular invasion (histological identification), respectively. The number of patients in each class is indicated. The fold change (LogFC) in gene expression is presented relative to mean expression level of the corresponding gene in five normal liver samples. Data are represented as Tukey’s boxplots where box indicates the first and third quartiles, bar indicates median, whiskers indicate 1.5 interquartile range (IQR). P-values obtained with the Wilcoxon rank-sum test are indicated. (**G**) Kaplan-Meier plots of overall survival in HCC patients (from(Nault et al., 2013)) with high or low expression of CDK8 or CDK19 mRNA. High/low expression groups were defined by the median expression levels of the total number of analyzed samples. Statistical differences were assessed by Log-rank Test. P-value obtained are indicated.

#### CDK8 is dispensable for liver homeostasis

To investigate possible roles of the Mediator kinases in liver function and hepatic carcinogenesis, we focused on CDK8. We generated a genetically modified mouse with loxP sites flanking exon 2 of the *Cdk8* gene (Figure 2A). Crossing these animals with transgenic mice expressing the *Cre* recombinase under the control of the *Albumin* promoter (*Alb-Cre* mice(29)) gives rise to hepatocyte-specific deletion of the essential exon 2, and induces a frameshift that results in a stop codon at position 52, eliminating CDK8 protein in the liver (CDK8^Δhep^ animals) (Figure 2B).

**Figure 2:**
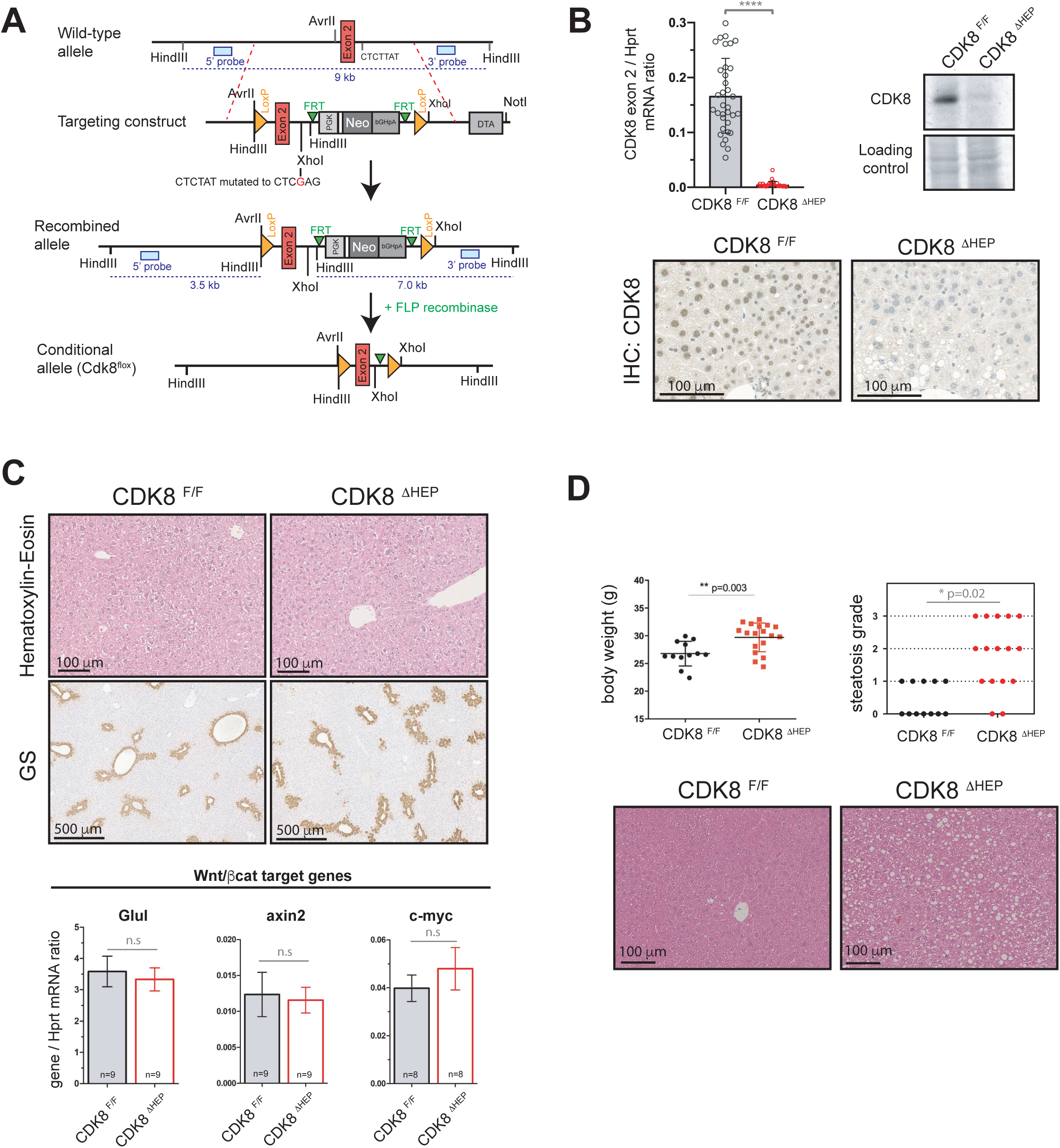
Hepato-specific ablation of CDK8 does not affect liver homeostasis, but induces steatosis. (**A**) Map of CDK8 genomic locus and strategy used to generate CDK8^F/F^ transgenic mouse. See material & methods section for details. (**B**) qPCR, Western-blot and immunohistochemical analysis of CDK8 removal in hepatocytes of CDK8^ΔHEP^ mice. For qPCR quantification, males and females mice of different ages were used. CDK8 ^F/F^ (n=34) or CDK8^ΔHEP^ (n=30). WB and IHC show results for 3 month old female mice. No differences were observed depending on sex or age of the animal. (**C**) HE staining of, GS staining, and qPCR analysis of β-catenin target genes (*Glul, axin2, c-myc*) in wild-type and CDK8 depleted 3 months old mice. n.s: not significant (t-test). (**D**) Body weight and steatosis grade score of 6 months old CDK8^F/F^ and CDK8^ΔHEP^ female mice. P value of Fisher’s exact test is indicated. Representative liver sections (HES staining) of mice from both genotypes are shown.

In accordance with the absence of overt phenotype of ubiquitous Cdk8 deletion in adult animals (10), the liver-specific deletion of CDK8 was compatible with normal liver development and we detected no change in liver physiology at the age of 3 months (Figure 2C). Importantly, the β-catenin-driven metabolic liver zonation was not affected by the CDK8 ablation, as judged by the purely centrilobular expression of glutamine synthetase (Figure 2C). Moreover, we detected no differences in the expression levels of several β-catenin target genes between the control and the CDK8^Δhep^ animals (Figure 2C). These results indicate that CDK8 is not required to regulate the β-catenin pathway under normal homeostasis conditions. However, in aging animals (> 6 months) CDK8 deficiency led to increased body weight and higher liver steatosis score (Figure 2D), confirming an involvement of CDK8 in liver lipogenesis (25). Older CDK8^Δhep^ animals (12-15 months) did not show any sign of liver tumors (n= 12).

#### CDK8 is required for chemically induced liver carcinogenesis and hepatic cell transformation

We next used a model of hepatic carcinogenesis in CDK8^Δhep^ animals to investigate potential roles of CDK8 in liver cancer. In this model, a single injection of hepatotoxic agent diethylnitrosamine (DEN) to young mice gives rise to liver tumors after 6-8 months (30). We sacrificed DEN-treated animals at 28 weeks, a relatively early time point in the kinetics of tumor formation, to allow detection of both positive and negative changes in tumor burden. As expected, 9 out 17 (53%) of the control *Cdk8*^*F/F*^ mice had at least one macroscopic liver tumor at sacrifice (Figure 3A). In contrast, only one out of eleven (9%) CDK8^Δhep^ animals had developed tumors by this time point. Cell death and regenerative response of the livers shortly after the DEN treatment were both indistinguishable between controls and CDK8^Δhep^ mice (Supplemental Figure 1), indicating that CDK8 ablation acted by inhibiting tumorigenesis rather than by interfering with the initial hepatoxicity of the treatment. Taken together, these data support the hypothesis that CDK8 contributes to chemically-induced liver carcinogenesis in mice.

**Figure 3:**
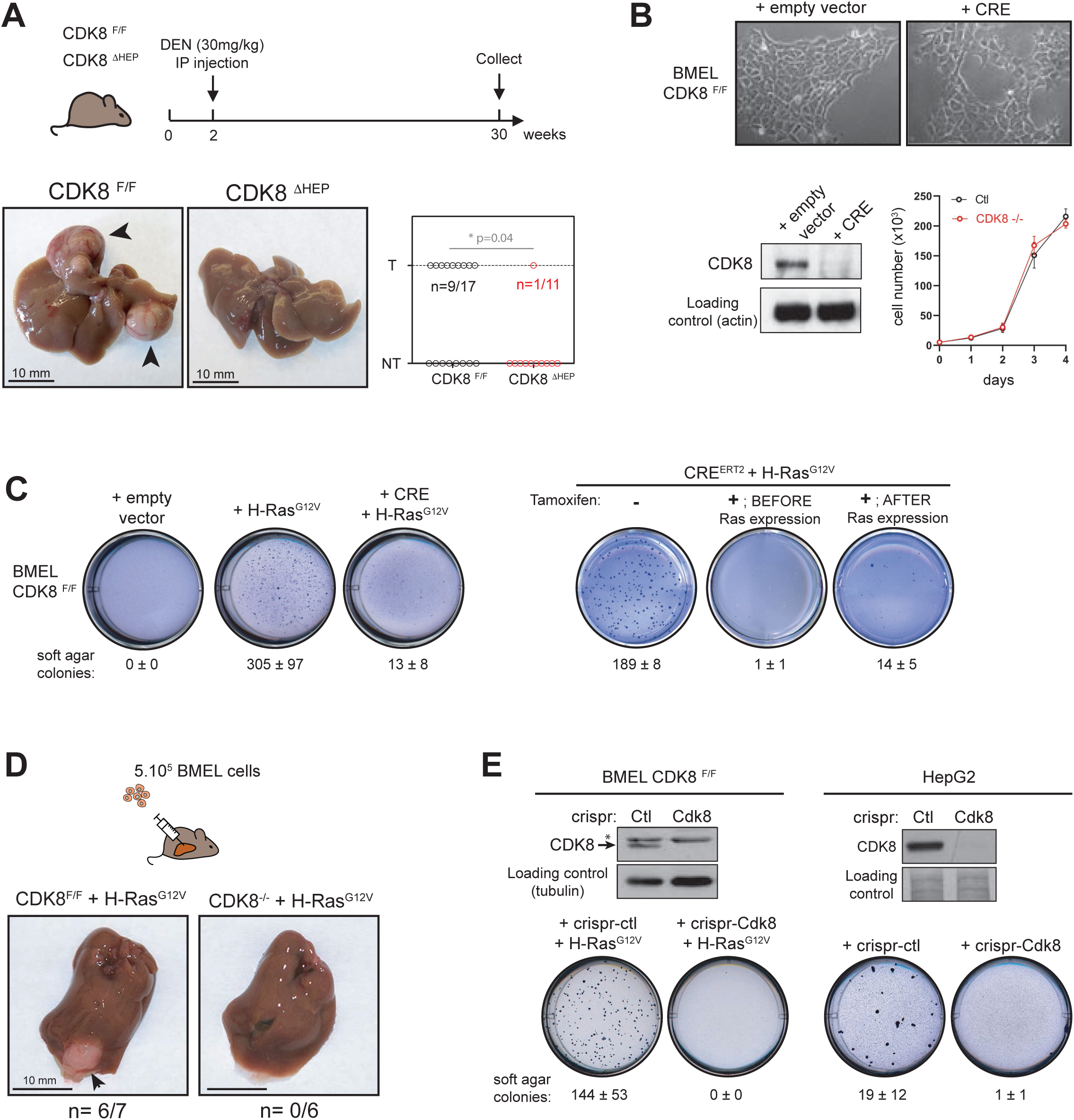
CDK8 is required for hepatic carcinogenesis. (**A**) Effect of CDK8 depletion on DEN-induced hepatic carcinogenesis. Pictures of representative livers for each genotype are shown. Arrowheads indicate tumors. Graph represents the repartition of the 28 DEN-injected animals, in each genotype depending if the liver presents a visible tumor (T) or not (NT). P-value of Fisher’s exact test is indicated. (**B**) Pictures and growth curve of CDK8^F/F^ hepatic progenitors (BMEL) WT or depleted of CDK8 by CRE recombination. (**C**) Soft agar growth of different BMEL cell lines. A representative well for each cell line is shown. Mean number of colonies per well ± SD from at least three independent experiments are indicated. Tamoxifen treatment (+Tam) was realized before or after transformation induced by H-Ras^G12V^ expression. (**D**) Result of orthotopic allografts of Ras expressing BMEL depleted or not of CDK8. A representative liver is shown, and the number of livers carrying a tumor out of total liver injected is indicated. Arrowhead points out to the tumor. (**E**) Western-blot characterization of CDK8 removal by crispr/cas9 gene editing and consequences on growth in soft agar for BMEL (left panel) and HepG2 (right panel) cells. Asterisk indicates an aspecific band sometimes detected in BMEL cells. Mean ± SD number of colonies per well from at least three independent experiments are indicated.

To investigate whether the effects of CDK8 loss on carcinogenesis were cell autonomous, we next isolated primary hepatic progenitor cells (BMEL) (31) from *Cdk8*^*F/F*^ embryos. Upon stable transfection of Cre, we obtained an efficient deletion of CDK8 from these cells (Figure 3B). CDK8 loss had no effect on BMEL cell morphology or growth characteristics under standard monolayer culture conditions (Figure 3B). As murine liver tumors triggered by DEN injection are often driven by mutated forms of Ras (30), we used forced expression of an oncogenic form of Ras, H-Ras^G12V^, to investigate effects of CDK8 loss on hepatic cell transformation. Similarly to previous results obtained with independent BMEL cell lines (32), H-Ras^G12V^ was sufficient to transform primary *Cdk8*^*F/F*^ BMEL cells, which then efficiently formed colonies in soft agar (Figure 3C). In contrast, CRE-mediated deletion of CDK8 abolished colony formation in this assay (Figure 3C), which is consistent with the protective role of CDK8 deletion in DEN-treated livers. Next, we used a model of tamoxifen-inducible Cre^ERT2^ activation. This confirmed the CDK8 requirement for Ras-induced transformation. Strikingly, deletion of CDK8 in cells previously transformed by Ras^G12V^ expression reverted their transformed phenotype (Figure 3C). To validate *in vivo* that CDK8 deletion protects hepatic progenitors from Ras-induced transformation, we injected CDK8^F/F^ Ras^G12V^ or CDK8^-/-^ Ras^G12V^ BMEL cells into the liver of immunodeficient mice. While the *Cdk8*^*F/F*^ BMEL expressing Ras^G12V^ gave rise to orthotopic tumors, their counterparts devoid of CDK8 did not (Figure 3D).

We next disrupted *Cdk8* by by CRISPR/Cas9-mediated gene disruption, wich confirmed both the absence of apparent phenotype in CDK8-depleted cells and the requirement for CDK8 for cell transformation (Figure 3E). Finally, to exclude that the requirement for CDK8 for transformation is specific for BMEL cells, we disrupted it by CRISPR/Cas9 in a human hepatoblastoma cell line, HepG2. As expected, the cells grew well in the absence of CDK8, but again did not form colonies in soft agar (Figure 3E). Altogether, our results indicate that CDK8 is required for H-Ras^G12V^-driven oncogenic transformation of hepatocytes and of hepatic progenitor cells.

#### CDK8 deletion impairs Ras^G12V^-driven transformation in a p53-dependent manner

We then investigated possible molecular mechanisms by which CDK8 removal could impair Ras^G12V^-driven transformation. Deletion of CDK8 did not abrogate Ras^G12V^-induced cell shape remodelling or ERK phosphorylation (Supplemental Figure 2A, B), indicating that the kinase is dispensable for Ras pathway activation. Second, constitutive activation of the β-catenin pathway in the BMEL cells did not rescue CDK8 deficiency (Supplemental Figure 2C), indicating that tumor-promoting activity of CDK8 in the liver does not rely on the activation of the β-catenin pathway, which is consistent with the lack of β-catenin-related phenotype in CDK8^Δhep^ livers (Figure 2). Third, CDK8 has been proposed as a regulator of glycolysis (33), and we therefore tested whether CDK8 removal affects the metabolism of BMEL cells. However, analysis of glycolysis (Seahorse Glycolysis stress test) and mitochondrial respiration (Seahorse Mito stress test) did not reveal any effect of CDK8 ablation (Supplemental Figure 3), indicating that this regulation is not present in hepatic progenitor cells and that it does not account for the failure of CDK8-mutant cells to grow in soft agar.

Since *CDK8* overexpression in patients correlates with mutant p53 status, we next considered the possibility that CDK8 acts by modulating p53 function. Although CDK8 can act as a coactivator of the p53 transcriptional program (34), we observed that removal of CDK8 rather increases the level of p53 protein (Figure 4A), suggesting that CDK8 might restrain the tumor-suppressive functions of p53 in hepatic cells. In agreement with this idea, soft agar tests showed that the requirement for CDK8 in Ras-induced transformation was abrogated by p53 inactivation via CRISPR/Cas9 editing (Figure 4B). Furthermore, depletion of CDK8 did not prevent transformation of the p53-deficient Huh7 cell line (Supplemental Figure 4). To extend these results to an *in vivo* setting, we triggered tumorigenesis via hydrodynamic gene delivery (HGD) (35) of the activated form of Ras together with CRISPR/Cas9-mediated inactivation of endogenous p53. HGD with Ras alone did not generate tumors, as previously described (36), whereas simultaneous transfection with N-Ras^G12D^ and CRISPR-p53 gave rise to multiple aggressive tumors within 4 weeks (Figure 4C). Consistent with the results of the cellular models, this combination of oncogenic stimuli was also fully efficient in CDK8^Δhep^ animals, whose hepatocytes are devoid of CDK8 (Figure 4C). Cell lines derived from CDK8^F/F^ or CDK8^-/-^ HGD-tumors were equally capable of giving rise to tumors when injected orthotopically into immunocompetent mice (Figure 4D). Our results suggest that CDK8 is required for initiation of tumorigenesis by counteracting p53 function.

**Figure 4:**
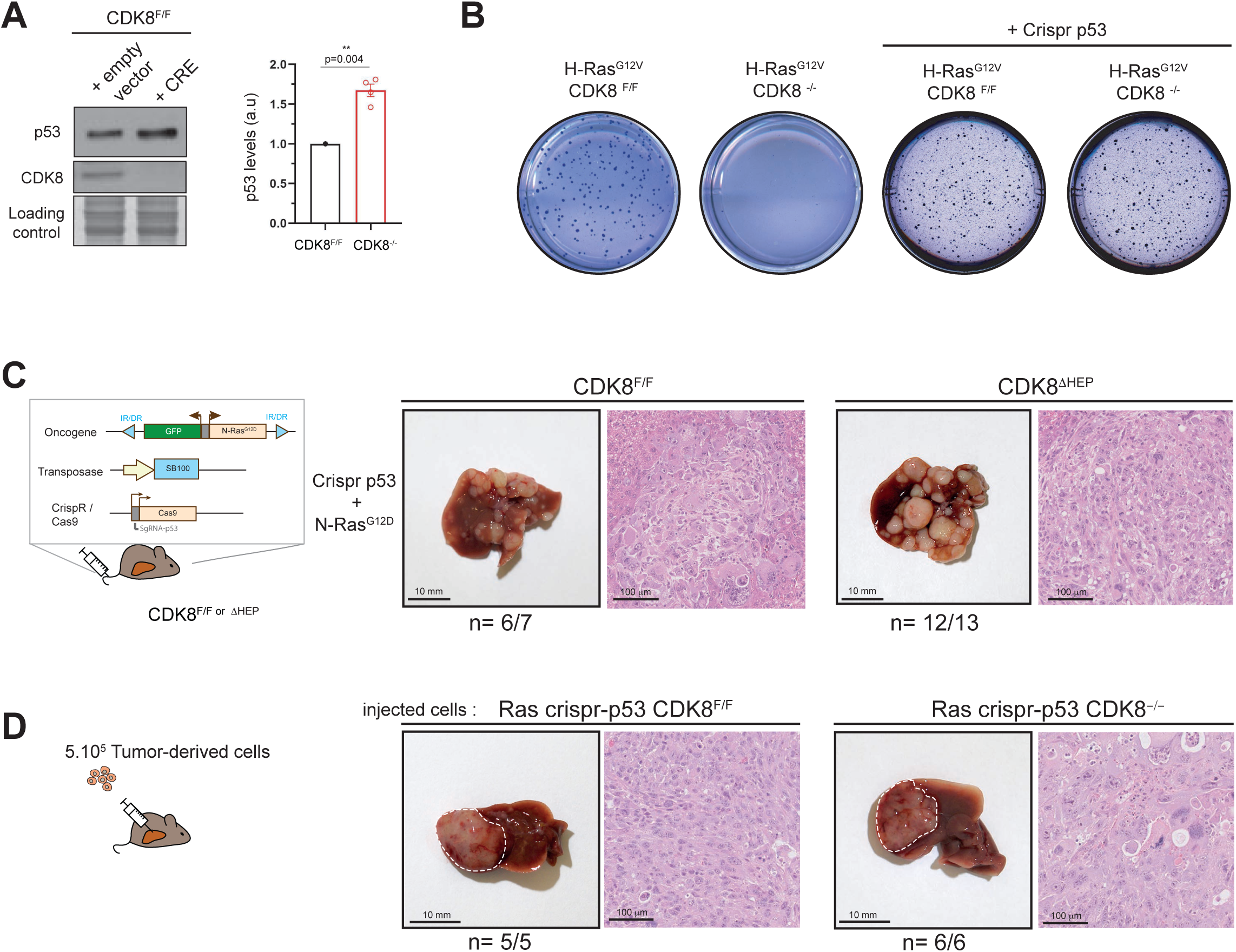
p53 deficiency abrogates CDK8 requirement for hepatic transformation. (**A**) Western Blot and quantification of p53 protein levels in BMEL cell lines, showing stabilization of p53 in absence of CDK8. A representative western blot and quantification from 4 independent experiments (mean ± SD) are shown. For each experiment, the band intensity value of the CDK8^F/F^ sample was normalized to 1. P-value from one-sample t test is indicated. (**B**) Soft agar test for indicated BMEL cell lines. (**C**) Hydrodynamic gene delivery of N-Ras^G12D^ and crispr-p53 vectors into livers of CDK8^F/F^ or CDK8^ΔHEP^ female mice. Representative HES staining of tumors are shown. For each genotype, number or mice presenting liver tumors out of number of mice injected is indicated. (**D**) orthotopic allografts of tumor cell lines derived from CDK8^F/F^ or CDK8^ΔHEP^ Ras crispr-p53. White dashed lines indicated tumor periphery. Representative HES staining of tumors are shown. For each genotype, number or mice presenting liver tumors out of number of mice injected is indicated.

#### Both CDK8 and CDK19 are required for Ras^G12V^-driven transformation

An interesting inference from the above results is that CDK19, whose expression is preserved in CDK8-deleted cells (Figure 5A), is insufficient to compensate for loss of CDK8, despite its high homology and redundant roles in Mediator. Furthermore, both CDK8 and CDK19 are overexpressed in a significant number of HCC patients. We therefore wondered whether CDK19 is also required for Ras-induced tumorigenesis. To test this, we deleted CDK19 in BMEL cells by CRISPR-Cas9 editing. The resulting mutant cells had unaltered morphology and proliferation kinetics (Figure 5A). Interestingly, CDK19 removal caused a significant upregulation of CDK8 protein levels, but not of mRNA levels, indicating a posttranscriptional feedback regulation of CDK8 in the absence of CDK19 (Figure 5A).

**Figure 5:**
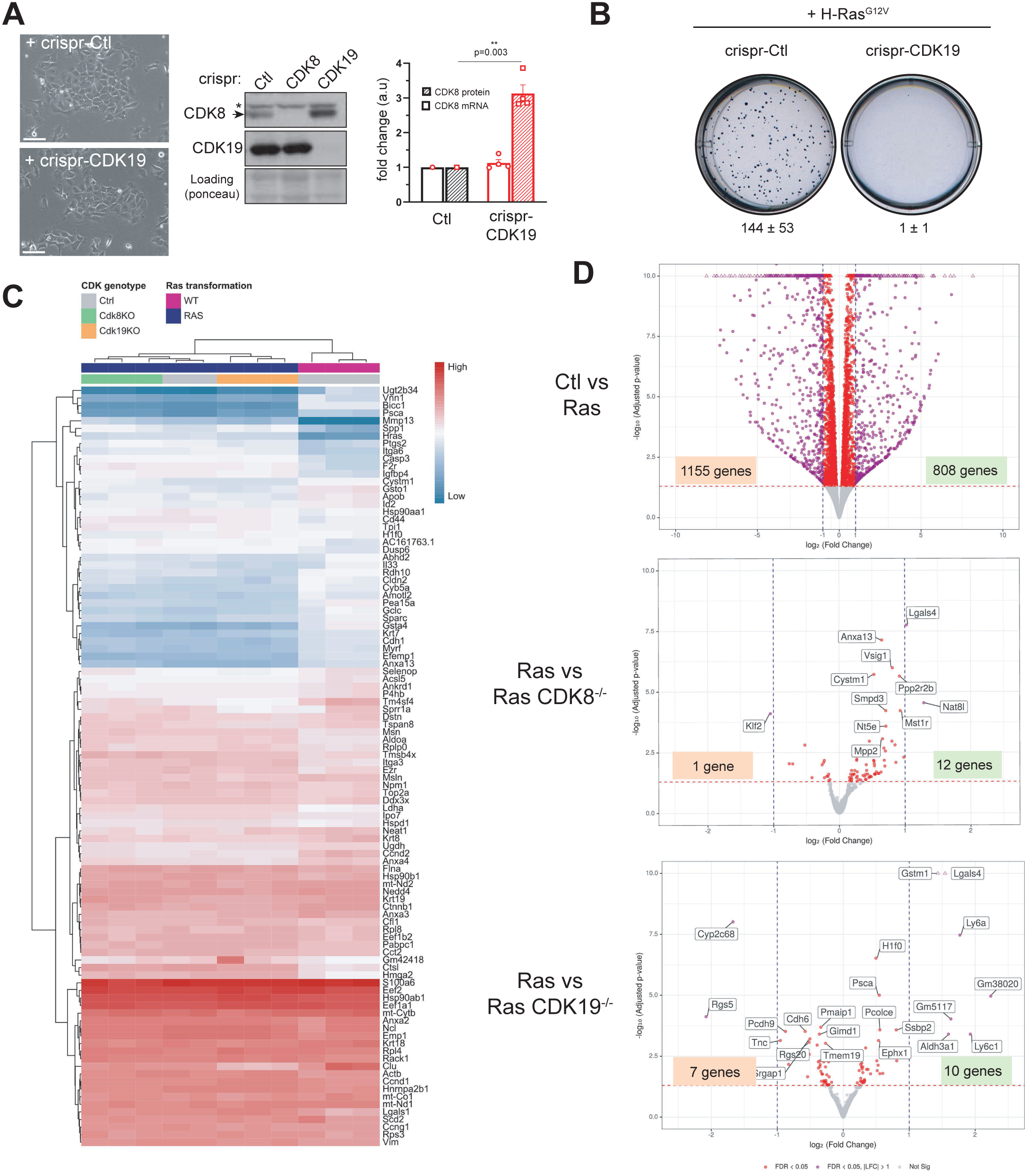
CDK19 and CDK8 are both required for transformation by H-Ras^G12V^. (**A**) Pictures, western blot and quantification of BMEL cell lines depleted of CDK19 by crispr/Cas9. A representative western blot and quantification from 4 independent experiments (mean ± SD) are shown. * indicates an aspecific band sometimes detected with CDK8 antibody. For each experiment, the band intensity value or cDNA content of the crispr-Ctl sample was normalized to 1. P-value from one-sample t test is indicated. (**B**) Soft agar test for indicated BMEL cell lines. A representative well for each cell line is shown. Mean number of colonies per well ± SD from at least three independent experiments are indicated. (**C**) Heatmap for the RNASeq experiments. The raw counts were transform using the regularized log and the variance was calculated between all the samples for all the genes. We selected the 100 genes with the higher variance. Distance between genes and samples were computed using the euclidean measure and the hierarchical cluster analysis was perform in order to set the distance trees. Two level of colors were set to describe the sample genotypes for RAS transformation (Violet: wild type; Blue: RAS^G12V^ mutation) and for Cdk8 or Cdk19 knock-outs (Gray: Control; Green: Cdk8 KO; Orange: Cdk19 KO). (**D**) Volcano plots for the differential gene expression. The values of the minus logarithm in base ten versus the logarithm in base two of the counts are plotted for each gene and two levels of significance are showed in red (genes with adjusted p-value less than 0.05) and in purple (genes with adjusted p-value less than 0.05 and with absolute values of the log2 of the fold change greater than one).

Similarly to CDK8 deletion, CDK19 ablation prevented the acquisition of the transformed phenotype upon ectopic expression of the oncogenic form of Ras, as judged by the lack of anchorage-independent growth (Figure 5B). Again, we obtained similar results upon CDK19 deletion in human HepG2 cells (Supplemental Figure 4). Thus, both CDK8 and CDK19 are required for Ras-driven transformation of primary hepatic progenitors, neither paralogue being able to compensate for the absence of the other.

As CDK8 and CDK19 constitute the only members of the CDK module regulating transcriptional activity of the mediator complex, one hypothesis to account for the requirement of simultaneous presence of both kinases is that they control different subsets of target genes, both of which are needed for transformation by H-Ras^G12V^. To test this hypothesis, we analyzed gene expression profiles of the Ras-expressing BMEL cells in the presence or absence of CDK8 or CDK19 by RNA-seq (Figure 5C). Surprisingly, disruption of either CDK8 or CDK19 had only minor effects on gene expression, with only around 30 genes reproducibly showing more than 2-fold differences in expression between cells expressing Ras^G12V^ with or without deletion of *Cdk8* or *Cdk19*. At this threshold of fold-change, we did not identify common genes deregulated by the absence of either kinase (Figure 5D). However, even though neither CDK8^-/-^ nor CDK19^-/-^ cells can be transformed by the oncogenic form of Ras, Ras^G12V^ caused deregulation of over 1000 genes, irrespective of the status of *Cdk8* and *Cdk19* when growing in exponential culture, explaining their indistinguishable morphology. This result further shows that, despite the importance of Mediator for gene regulation, neither CDK8 nor CDK19 are required for implementing wide-ranging changes to gene expression, suggesting that their effects in carcinogenesis, which involve p53, are not due to major transcriptional changes.

## Discussion

Combining patient data and mouse models, we provide evidence that both Mediator kinases, CDK8 and CDK19, are involved in hepatocellular carcinoma. Our data indicate that these CDKs are required for initiation of tumorigenesis induced by mutation of a strong oncogene, *Ras*. To our knowledge, this is the first indication that the CDK8 paralogue CDK19 is required for cell transformation, indicating non-redundant functions for both kinases in facilitating tumorigenesis.

We find that removal of CDK8 in hepatocytes has no major consequences for liver functions, as previously indicated by the ubiquitous inactivation of the kinase in adult animals (10). However, we observed that CDK8 depletion induces steatosis in aging animals, in accordance with a role for the kinase as repressor of lipogenesis (25). In contrast to its mild effects on liver physiology, CDK8 disruption has a strong impact on hepatic carcinogenesis. We observed a major reduction of chemically induced tumorigenesis, and a complete protection from Ras-induced cell transformation. This requirement is further highlighted by the fact that CDK8 ablation occurring after Ras^G12V^-induced transformation is able to restore the non-transformed phenotype.

CDK8 thus apparently has diverse context-specific involvement in various cancers. CDK8 was initially described to have oncogenic properties in Wnt-dependent colorectal cancers, where it controls the beta-catenin pathway (6, 8) and maintains Myc functions (37). However, this was not confirmed genetically in mouse models; indeed, if anything, CDK8 knockout marginally increased progression from early lesions to tumors (10). A recent study also found no effect of CDK8 knockdown on growth of colorectal tumors in syngeneic mice, but it prevented liver metastases (38). Furthermore, CDK8 promotes proliferation of melanoma cell lines (7) and is apparently required for efficient Estrogen Receptor (ER)-dependent transcription in ER-positive breast cancer cells (39). However, most reports of anti-tumor activity caused by interfering with CDK8 function have used kinase inhibitors (39–45) that target both CDK8 and CDK19. The most likely explanation for this is that CDK inhibitors are rarely very specific for one kinase subfamily (46) and some “CDK8/19 inhibitors” may inhibit other kinases required for efficient tumor development. This highlights the need for a genetic approach *in vivo*.

Other studies have suggested that as well as possessing oncogenic activity in some contexts, CDK8 may rather have tumor-suppressive activity in others, including endometrial cancers (47), intestinal cancer (10) and T-cell acute lymphoblastic leukemia (3).

One way of reconciling these apparently different roles for Mediator kinases is to invoke possible tissue-specificity of action. We found that CDK8 deletion has no effect on hepatic cell growth, contrary to certain melanoma or gastric cancer cell lines, in which decrease of proliferation has been described (7, 48). Tissue specificity is further demonstrated by the fact that while knockout of either kinase is well tolerated by hepatic precursor cells, germline-deletion of the *Cdk8* gene in mice is lethal at pre-implantation embryogenic stages (2). Our results suggest that oncogenic activities of CDK8 and CDK19 in HCC are independent of the β-catenin pathway, and therefore likely operate through different mechanisms in colorectal and hepatic carcinogenesis. One key finding of our study is that both CDK8 and CDK19 are required for cell transformation by H-Ras^G12V^. This is not due to downregulation of one kinase when the other one is removed, as, on the contrary, we found that CDK19 depletion stabilizes CDK8 protein, identifying a post transcriptional regulation of CDK8 levels. Of note, this cross-regulation is not bidirectional, as removal of CDK8 does not seem to stabilize CDK19 protein. A possible explanation for these results might be that CDK8 and CDK19 regulate transcription of specific gene sets that are both required for hepatic cell transformation. The results of our RNA-seq analysis renders this hypothesis unlikely, as very few genes are differentially transcribed in either knockout. However, we cannot exclude this entirely, as 12 genes were differentially regulated in knockouts of both *Cdk8* and *Cdk19* when the threshold for fold-change is removed: Adora1, Nt5e, Fgfbp1, Spp1, Gm5781, Lgals4, Rpl10-ps3, Akap12, Gm8349, Anxa10, Scd1, Ly6a. Yet the magnitude of the gene expression change is likely too low to be biologically meaningful; the regulation of several, including Anxa10 (encoding Annexin 10A), Akap12 (encoding A Kinase Anchoring Protein-12) and Nt5e (encoding 5’ nucleotidase) is in opposite directions in each knockout; and no clear roles in cancer for any of these genes have been reported. A possibility that we cannot exclude at this stage is that both CDK8 and CDK19 are required for transcription or repression of genes whose expression is only activated upon growth in foci. However, the minimal effects of deletion of either gene in shaping the transcriptome induced by expression of oncogenic Ras suggests that neither kinase is required to implement large-scale changes to gene expression, and that non-transcriptome effects should be considered.

We suggest an alternative model, which is compatible with both apparent oncogenic activity of Mediator kinases in some circumstances and lack of effects in others. In this model, CDK8/19 are not in themselves oncogenes and therefore their overexpression does not transform cells, but rather provides a favorable terrain enabling the initiation of tumorigenesis by additional oncogenic factors, including *bona fide* oncogenes. For example, it is well established that, rather than transforming cells directly, expression of strong oncogenes such as Ras^G12V^ promotes premature cell senescence with accumulation of p53 and p16 (49). BMEL cells are immortal, but have a functional wild-type p53 (50). In this context CDK8/19 expression may attenuate the function of p53, thus unleashing the oncogenic potential of mutant Ras. We speculate that loss of p53 by mutation “fixes” the initial advantage conferred by overexpression of CDK8/19. This agrees both with our finding that in the liver, CDK8/19 are only required for cell transformation in the presence, but not the absence, of p53, and with the genetic interaction between *CDK8/19* and *TP53* in patients. If such a scenario was indeed true, it would suggest a therapeutic niche for CDK8/19 kinase inhibitors in cancer: they would be expected to trigger differentiation or death of tumor cells that have not yet acquired p53 mutations. Our results warrant further investigation of CDK8/19 inhibitors as therapeutic agents for p53-positive hepatocellular carcinoma.

## Material and Methods

### Patients

A total of 268 fresh-frozen tissue samples of HCC, associated with various etiologies, were included in this study. Patients and tumor features were already described in previously published studies and summarized in Supplemental Table 1. Written informed consent was obtained from all subjects in accordance with French legislation.

*CDK18* and *CDK19* mRNA expression levels were assessed by quantitative RT-PCR using Fluidigm 96.96 Dynamic Arrays and specific TaqMan predesigned assays (CDK8= Hs00176209_m1; CDK19= Hs00292369_m1; Life Technologies, Carlsbad, CA). Data were calibrated with the RNA ribosomal 18S and changes in mRNA expression levels were determined using a comparative CT method using 5 normal tissue samples as control.

### Mice experiments

All reported animal procedures were carried out in accordance with the rules of the French Institutional Animal Care and Use Committee and European Community Council (2010/63/EU). Animal studies were approved by institutional ethical committee (Comité d’éthique en expérimentation animale Languedoc-Roussillon (#36)) and by the Ministère de l’Enseignement Supérieur, de la Recherche et de l’Innovation (D. Gregoire: APAFIS#11196-2018090515538313v2).

Cdk8^F/F^ mice were generated as following: An 8076 bp genomic fragment (mouse chromosome 5: 146,254,503 to 146,262,579) enclosing the essential exon 2 (whose deletion results in loss of the essential catalytic lysine residue and causes a frameshift truncating over 90% of the protein) of the CDK8 gene was amplified by PCR from genomic DNA of 129/Sv embryonic stem cells and cloned into pGEM-T-easy. The diphtheria toxin A gene was cloned into the SacII site. 64 bp to the 3’ of exon 2, the sequence CTCTAT was mutated to CTCGAG, generating an XhoI site. LoxP sites flanking exon 2 were generated by a combination of conventional cloning and recombineering, using an approach published in Liu et al., Genome Res. 2003 Mar;13(3):476-84. The loxP PGK-Neo cassette was amplified from pL452 plasmid with flanking AvrII/HindIII sites at each end and cloned into the AvrII site upstream of exon 2. Fragment orientation was confirmed by the generation of 3.5 kb HindII and 2.0 kb NheI sites, and the vector was recombined in *E*.*coli* strain SW106 with inducible Cre recombinase expression followed by HindII digestion, generating a single loxP site upstream of exon 2. Into this recombined vector, the FRT-PGK-Neo-FRT-LoxP cassette (amplified from pL451 with flanking XhoI sites) was cloned in the newly generated XhoI site downstream of exon 2, resulting in the “deletion construct”. The orientation was confirmed by the generation of 2.2 kb Nhe1 and 3.4 kb BamHI sites. Functionality of the two recombination sites was tested as follows: the FRT site was confirmed by recombination in *E*.*coli* strain SW105 with inducible FlpE recombinase expression, deleting the FRT-Neo cassette and generating a 1.4 kb BamHI fragment; and the resulting plasmid was transformed in *E*.*coli* strain SW106 with inducible Cre recombinase expression, deleting exon 2 and resulting in a 1.1 kb BamHI fragment. The NotI linearized fragment of the deletion construct was transfected by electroporation into 129/Sv embryonic stem cells. 244 Neomycin-resistant colonies were genotyped by PCR and Southern blotting. Two probes were used: one outside the 3’ end of the deletion construct, with HindIII digestion giving a single 9kb fragment for the WT and a 7kb fragment for the correctly-integrated deletion cassette, and one to the 5’ end of the deletion cassette, again giving the same 9kb fragment for the WT but a 3.5 kb fragment for the deletion cassette. 10 colonies showed a correct integration by homologous recombination. These ES cells were injected into blastocysts obtained from pregnant Balb/c mice, and chimeric mice were crossed with C57/Bl6J mice constitutively expressing FlpE recombinase, removing the FRT-Neo cassette. Agouti mice were genotyped by PCR, showing correct insertion of the loxP sites around exon 2. Alb-Cre mice were described previously (29).

Allografts: Athymic Nude mice (*Hsd:Athymic Nude-Foxn1*^*nu*^, Envigo) or CDK8^F/F^ mice were anesthetized with intra-peritoneal injection of Xylazine-Ketamine mixture. After incision of abdominal wall and peritoneum, the left lateral lobe of the liver was pulled out of the mouse body. 50000 cells, resuspended in 5 μL of 25% Matrigel (BD) - PBS, were injected using a 10μL Hamilton syringe in the left lobe of the liver. After injection, liver was put back in normal position and the abdomen was sutured. Tumors were allowed to grow out for 4 weeks, then collected and fixed following classical procedures.

#### DEN induced carcinogenesis

Diethylnitrosamine (DEN) (30 mg/kg) was injected intra-peritoneally in 14 days old male mice. Mice were sacrificed and livers collected after a period of 8 months.

#### Hydrodynamic Gene Delivery

Hydrodynamic injections were performed in 6-8 week-old female mice as described previously (35). Briefly, 0.1 mL/g of a solution of sterile saline (0.9% NaCl) containing plasmids of interest were injected into lateral tail vein in 8-10 s. LentiCRISPRv2-sgTp53 (12.5 µg) and pT3-EF1a-N-RAS^G12D^-GFP (12.5 µg) were injected together with sleeping beauty transposase SB100X (2.5 µg, ratio of 5:1). pCMV(CAT)T7-SB100 was a gift from Zsuzsanna Izsvak (Addgene plasmid # 34879).

### Cell lines

BMEL (Bipotential Mouse Embryonic Liver) cell line was isolated from CDK8^F/F^ mouse. Cells were grown on collagen-coated plates in RPMI medium (Gibco/Life Technologies) supplemented with 10% fetal calf serum (Pan-Biotech), insulin 10 μg/mL (Sigma), IGFII 30 ng/mL (Peprotech), EGF 50 ng/mL (Peprotech), 100 units/ml penicillin, and 100 mg/ml streptomycin. Other cell lines used, HepG2, Huh7, and HEK293T, were grown in Dulbecco modified Eagle medium (DMEM–high glucose, pyruvate, GlutaMAX–Gibco^®^ LifeTechnologies) supplemented with 10% fetal bovine serum (Pan-Biotech). Cells were grown under standard conditions at 37°C in a humidified incubator containing 5% CO_2_. All cells were routinely tested to confirm absence of mycoplasma contamination.

### Generation of cell lines

pMSCV retroviral vectors (Clontech) encoding CRE recombinase, tamoxifen-inducible CRE-ERT2 (addgene plasmid #22776) or human H-Ras^G12V^ were used to generate stable BMEL cell lines. For induction of CRE-ERT2 activity, 4-OH tamoxifen was added to culture medium at a concentration of 2 μM for two weeks. CRISPR subgenomic RNA (sgRNA) targeting murine or human Cdk8, Cdk19 or Trp53 (CDK8 Mouse: 5’-ATCCCTGTGCAACACCCAGT-3’. CDK8_Human 5’-CGAGGACCTGTTTGAATACG-3’ CDK19_Mouse: 5’-AAAGTGGGACGCGGCACCTA-3’, CDK19_Human, 5’-ATTATGCAGAGCATGACTTG-3’ Trp53 5’-ATAAGCCTGAAAATGTCTCC-3’; all sequences from Zhang lab database) were cloned as synthetic dsDNA into lentiCRISPRv2 vector as described (51) (provided by F. Zhang, Addgene plasmid #52961). Generation of lentiviral particles and infection of BMEL, Huh7 or HepG2 cells were carried out following classical procedures, and described previously (52). Successfully infected cells were selected with puromycin (2 μg/mL) or hygromycin (150 μg/mL) during 48h. Cell lines were further propagated and transgene expression or effects of targeted deletions verified by Western blot. Polyclonal cell lines were used for all subsequent experiments.

### Soft agar

10^5^ cells for each cell line were mixed with medium supplemented with 0.5% agarose and placed on top of the 1% agarose layer. 1 mL medium was added to the solidified layer and changed every 2-3 days. After 4 to 6 weeks, soft agar was stained with crystal violet 0,005% in 4% PFA for 1h. Colonies visible to the naked eye were counted manually.

### Immunohistochemistry

Livers were fixed for 24 h in 10% neutral buffered formalin, embedded in paraffin, sectioned at 4 μm and stained with haematoxylin-eosin safran (HES) or subjected to immunohistochemical staining. Immunohistochemistry staining of GS (BD Transduction Laboratories, 610517, 1:200) and CDK8 (Santa Cruz, sc-1521, 1:400) were performed using classical procedures, with antigen retrieval in citrate buffer, and biotinylated secondary antibody coupled to streptavidin– peroxidase complex (ABC Vectastain kit; Vector Laboratories). Slides were digitally processed using the Nanozoomer scanner (Hamamatsu).

### Protein isolation and Western blotting

Protein lysates of cells or tissues were prepared with lysis buffer (150mM NaCl, 50mM Tris pH 7.5, 0.2% Triton, 1mM EDTA; freshly added 1mM DTT and protease inhibitors cocktail (Roche)) and incubated on ice for 30 min. Samples were centrifuged for 10 min at 13000rpm and supernatant collected. Protein concentrations were determined by BCA protein assay (Pierce Biotechnology). All the samples were mixed 1:1 with Laemmli buffer and heated at 95°C for 5 min. Equal amounts of proteins were separated by SDS–PAGE (Biorad; usually 12% gels). Proteins were transferred onto PVDF membranes (Milipore). Red ponceau or amidoblack staining was done to check transfer efficiency and equal amount of protein loading. Primary antibodies used were: Cdk8 (Santa Cruz #sc1521, dilution 1:1000); Cdk19 (Abcam, dilution 1:1000); p53 (Cell signalling #2524 1:1000), Actin (Sigma A1978, 1:20000), Tubulin (DSHB, 1:400). Antibodies were diluted in 5% BSA in TBS-Tween and incubated overnight at 4°C. Secondary antibodies were either anti-goat IgG-HRP or anti-mouse antibodies IgG-HRP (Jackson immunoresearch). Band intensities were calculated using imageJ Lane Analysis.

### RNA isolation, qPCR and RNA-Seq analysis

The RNA was extracted from either cells or liver tissue and purified using RNeasy mini kit (Qiagen) according to manufacturer’s protocol. Reverse transcription of total RNA (1μg) was done with QuantiTect Reverse Transcription kit (Qiagen), and cDNA quantified using LC Fast stard DNA Master SYBR Green I Mix (Roche) with primers detailed below on LightCycler480 apparatus (Roche). Gene expression levels were normalized with hypoxanthine phosphoribosyltransferase (HPRT). Primer pairs used for qPCR: Cdk8 5’-GAATTTCTATGTCGGCATGCAG-3’ and 5’-ATAGTCAAAGAGAAGCCATACTTTCC-3’, Glul 5’-TAGCTGTCACAAAGCGGGTGTA-3’ and 5’-AGTGGAAATGTCAATCTCAGCC-3’, Axin2 5’-ACCGGTCACAGGATGTC-3’ and 5’-GACTCCAATGGGTAGCTCTTTC-3’, c-myc 5’-CCGAGTGCATTGACCCCTCA-3’ and 5’-GAGAAGGCCGTGGAATCGGA-3’. Hprt 5’-GCAGTACAGCCCCAAAATGG-3’ and 5’-GGTCCTTTTCACCAGCAAGCT-3’.

For RNA-Seq analysis, RNA was extracted from exponentially growing subconfluent BMEL cells in three independent experiments for each cell line, using RNeasy mini kit (Qiagen) with DNAse treatment. RNA integrity was validated using RNA BioAnalyzer (Agilent), all RIN ≥ 9.5. The preparation of the library was done with the TruSeq Stranded mRNA Sample Preparation kit (Illumina). The sequencing was performed in an Illumina Hiseq 2500 sequencer by the Sequencing Platform of Montpellier (GenomiX, MGX, France; www.mgx.cnrs.fr), with 50 base pairs (bp) single end reads to an estimated depth of 25 million reads per sample. In order to perform a quality control of the sequencing, FastQC over the fastq files containing the raw reads. All the reads that passed the quality control were aligned to the mouse reference genome (GRCm38.p6) and the counts per gene were quantified using the tool STAR 2.6.0a(2). The Ensembl mouse genome annotations (release 93) were used for establishing the coordinates of each gene and their corresponding transcripts. Differential gene expression analysis was performed in R using the DESeq2 library. After normalization of the read counts per gene, a negative binomial generalized linear model was fitted considering single factor design for assessing the differential expression between CDK8/CDK19 knock-out BMEL Ras^G12V^ transformed cells and BMEL Ras^G12V^ transformed cells (as the control group). Wald test are performed for assessing statistical significance on the differential expression of each gene, then test are independently filtered and corrected by multiple hypothesis testing (Benjamini–Hochberg).

### Statistical Analysis

Data sets were tested with 2-tailed unpaired Student *t* tests or Mann-Whitney U tests, correlations were analyzed with Pearson’s χ^2^ test using Prism Software version 8 (GraphPad). Significant *P* values are shown as: \**P* <0.05, \*\**P* <0.01, \*\*\**P* <0.001, and \*\*\*\**P* <0.0001.

Human samples: Data visualization and statistical analysis were performed using R software version 3.5.1 (R Foundation for Statistical Computing, Vienna, Austria. https://www.R-project.org) and Bioconductor packages. Comparisons of the mRNA expression levels between groups were assessed using Mann-Whitney U test. Spearman’s rank-order correlation was used to test the association between continuous variables. Univariate survival analysis was performed using Kaplan-Meier curve with log-rank test. The median *CDK8* and *CDK19* expression levels on the total number of analyzed samples was used to determine the low- and high-expression groups. P-value < 0.05 was considered as significant.

## Data availability

The RNA-sequencing data have been deposited in the Gene Expression Omnibus (GEO, NCBI) repository.

## Author contributions

KB, SP, DG performed and analyzed experiments. SC and JZR acquired and analyzed human patients data. AC generated CDK8^F/F^ transgenic mouse line. GD analyzed RNA-Seq data. JUB, AL, JB contributed to *in vivo* experiments. DG made the figures. UH, DF and DG designed experiments, analyzed data, supervised the study and wrote the manuscript.

### Acknowledgments

We acknowledge Montpellier Biocampus facilities: the imaging facility (MRI), the “Réseau d’Histologie Expérimentale de Montpellier” (RHEM) and Montpellier Genomix (MGX). We are grateful to zootechnicians of IGMM animal housing facility for their work. We thank Christina Begon-Pescia (IGMM transgenesis facility) for help in generation of CDK8 transgenic mouse line. We thank Scott Lowe for the gift of pT3-EF1a-N-RAS^G12D^-GFP plasmid and Leila Akkari for help setting up hydrodynamic injections. We thank members of our labs for helpful discussions and comments.

This work was funded by Institut National du Cancer (INCa; PLBIO 2015-132, DF and UH teams) and EVA-Plan cancer INSERM THE (UH, JZR), and supported by SIRIC Montpellier Cancer Grant INCa_Inserm_DGOS_12553. DF is funded by the Ligue Nationale contre le Cancer (EL2017-LNCC/DF). DG benefited from support by Association Française pour l’Etude du Foie (AFEF). JZR team is supported by the Ligue Nationale Contre le Cancer (Equipe Labellisée), Labex OncoImmunology (Investissement d’Avenir), the Fondation Bettencourt-Schueller (coup d’élan Award), the Ligue Contre le Cancer Comité de Paris (Duquesne award) and the Fondation pour la Recherche Médicale (Raymond Rosen award). SC is supported by a funding from “Labex OncoImunology”, KB by INCa;PLBIO and JB by ANRS. The funders had no role in study design, data collection and analysis or publication process.

## ID ORCID

Susana Prieto https://orcid.org/000-002-9746-2396

José Ursic-Bedoya: https://orcid.org/0000-0003-0076-2059

Stefano Caruso: https://orcid.org/0000-0002-6319-3642

Anthony Lozano: https://orcid.org/0000-0002-9749-0573

Jessica Zucman-Rossi: https://orcid.org/0000-0002-5687-0334

Urszula Hibner: https://orcid.org/0000-0002-5520-7311

Daniel Fisher: https://orcid.org/0000-0002-0822-3482

Damien Gregoire: https://orcid.org/0000-0002-1105-8115

