## Supplemental figures for "CDK8 and CDK19 kinases have non-redundant oncogenic functions in hepatocellular carcinoma"

**Supplemental Table 1. Clinical and Molecular Characteristics of the HCC samples (n = 268).**

| Variable | | Total (%) |
|---|---|---|
| **Age (n=268)** (years) | [median, min-max] | 65 (18-90) |
| **Gender** | Male : Female | 221 (82%) : 47 (18%) |
| **Etiology** (including 59 patients with ≥ 2 etiologies) | Alcohol | 115 (43%) |
| | Hepatitis B | 55 (20.5%) |
| | Hepatitis C | 56 (21%) |
| | Hemochromatosis | 23 (9%) |
| | Metabolic syndrome | 44 (16%) |
| | Other etiologies | 1 (0.5%) |
| | Without known etiology | 34 (13%) |
| **Edmonson grading (n=265)** | I | 8 (3%) |
| | II | 114 (43%) |
| | III | 117 (44%) |
| | IV | 26 (10%) |
| **Differentiation WHO (n=267)** | Good | 69 (26%) |
| | Medium | 156 (58%) |
| | Week | 42 (16%) |
| **G1G6 (n=268)** (Boyault S. et al., Hepatology 2007) | G1 | 16 (6%) |
| | G2 | 23 (9%) |
| | G3 | 47 (17%) |
| | G4 | 87 (32%) |
| | G5 | 56 (21%) |
| | G6 | 39 (15%) |
| **5-gene score (n=257)** (Nault JC et al., Gastroenterology 2013) | P1 | 142 (55%) |
| | P2 | 115 (45%) |
| ***TERT* promoter mutations (n=252)** | Mutated | 160 (63%) |
| ***CTNNB1* gene mutations (n=258)** | Mutated | 106 (41%) |
| ***TP53* gene mutations (n=263)** | Mutated | 54 (21%) |

#### Supplemental Figure 1_Bacevic et al.

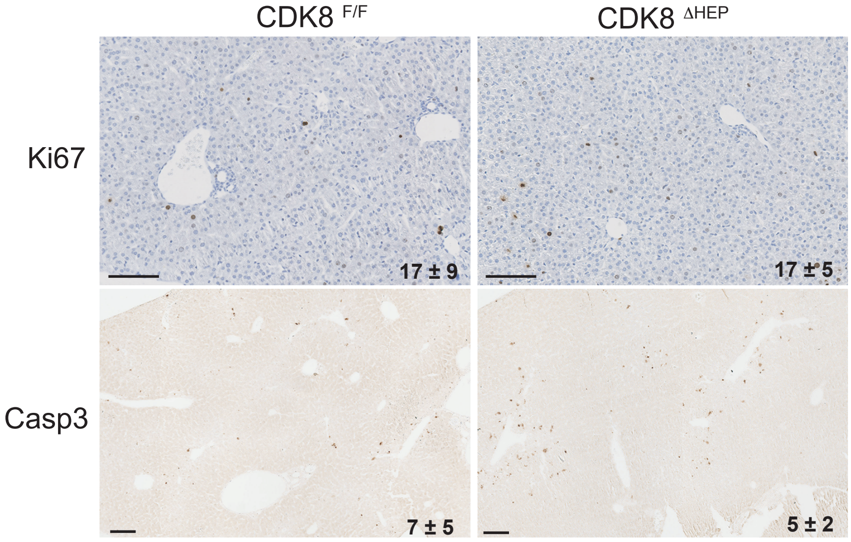


**SuppFigure 1: CDK8 ablation does not affect response to DEN injection.** Immunohistochemistry of Ki67 (cell proliferation) and activated caspase 3 (apoptosis) livers 48h after intraperitoneal injection of DEN (30 mg/kg) in $CDK8^{F/F}$ and $CDK8^{\Delta HEP}$ two-weeks old males. Mean ± SD number of positive cells per field are indicated. Scales bar: 100 μm

### Supplemental Figure 2 _Bacevic et al.

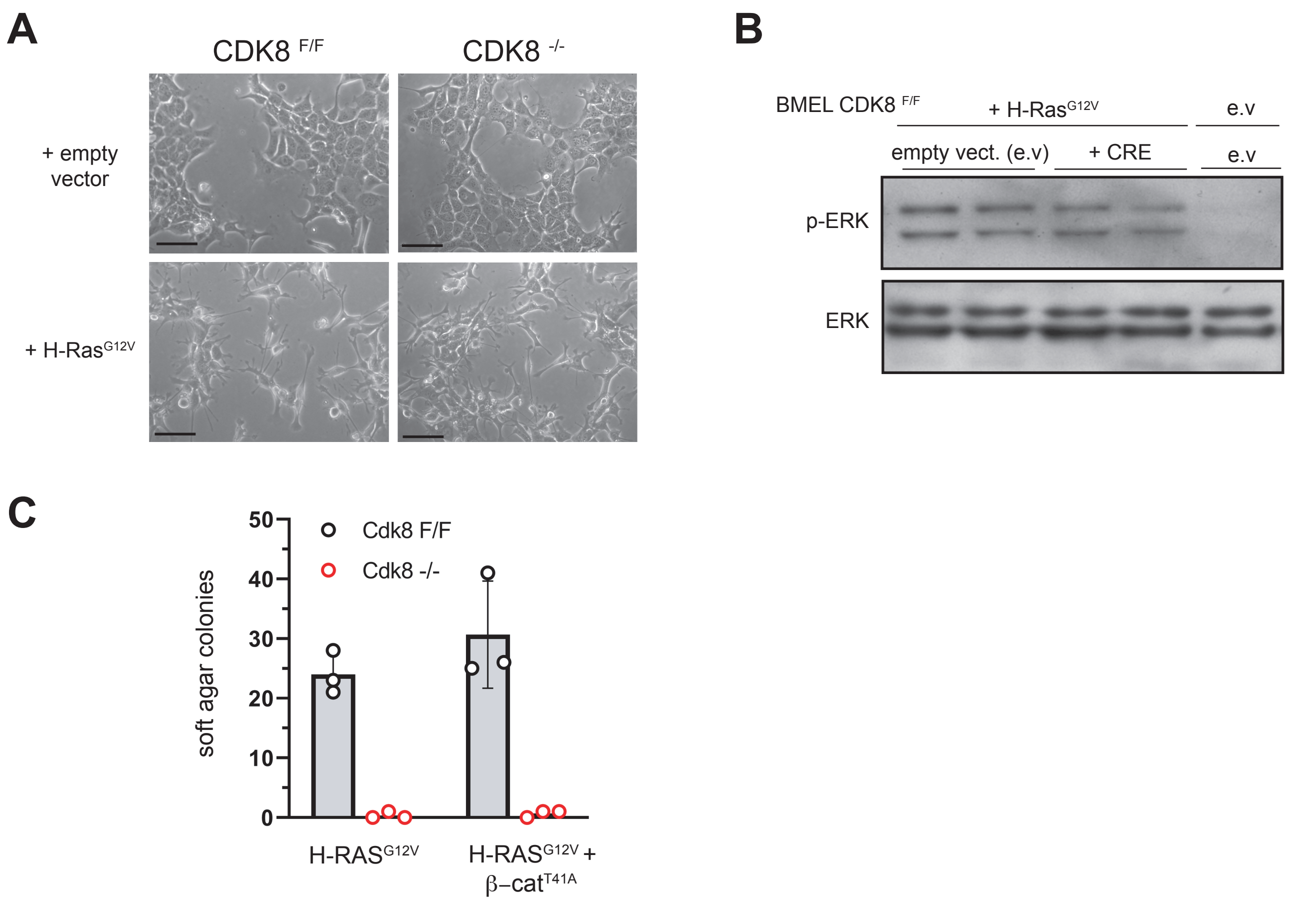


**SuppFigure 2: CDK8 ablation impact on cell transformation does not depend on Ras or β−catenin pathways.** (**A**) phase contrast microscopy of subconfluent CDK8$^{F/F}$ BMEL cells expressing or not Cre recombinase and H-Ras$^{G12V}$. (**B**) Western blot analysis of ERK phosporylation subconfluent CDK8$^{F/F}$ BMEL cells expressing or not Cre recombinase and H-Ras$^{G12V}$. e.v= empty vector, pMSCV with no insert. (**C**) Number of colonies growing in soft agar conditions for different BMEL cell lines. Mean ± SD number of colonies from three different wells of a representative experiment are shown. Note that expression of active β-catenin does not rescue growth capacities in CDK8$^{-/-}$ BMEL cells.

#### Supplemental Figure 3 _ Bacevic et al.

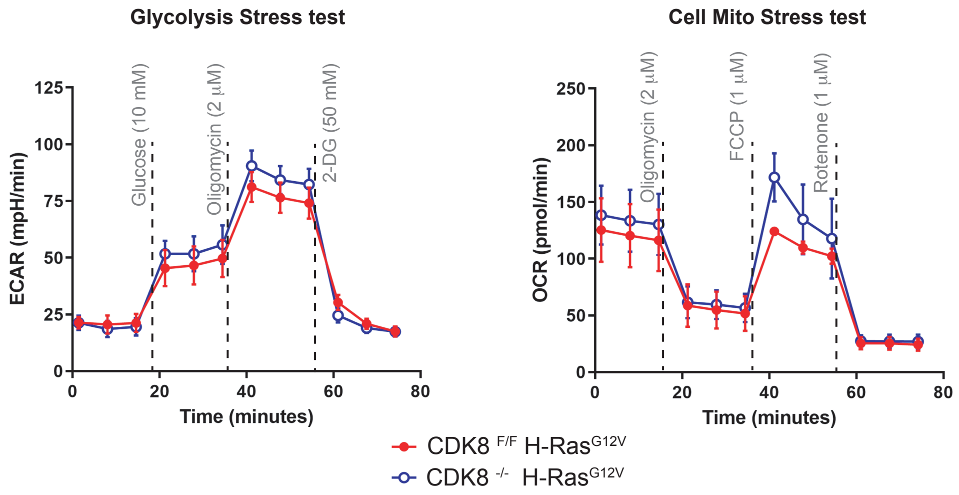


**SuppFigure 3: CDK8 removal in BMEL does not affect glycolysis or mitochondrial respiration.** Results of seahorse glycolysis stress test (Agilent) and Cell Mito stress test (Agilent) for BMEL expressing H-Ras$^{G12V}$ in which CDK8 has been depleted by Cre recombinase (CDK8$^{-/-}$) or not (CDK8$^{F/F}$). ECAR: ExtraCellular Acidification Rate; OCR: Oxygen Consumption Rate. Mean ± SEM of quadruplicates are shown. Tests were carried out on a Seahorse XFe96 (Agilent), using Glycolysis Stress Kit (Agilent) and Cell Mito Test Kit (Agilent), following manufacturer's instructions.These results were reproduced in an independant experiment and similar results were obtained with cell lines in which CDK8 was removed by crispr/Cas9 editing.

#### Supplemental Figure 4 _ Bacevic et al.

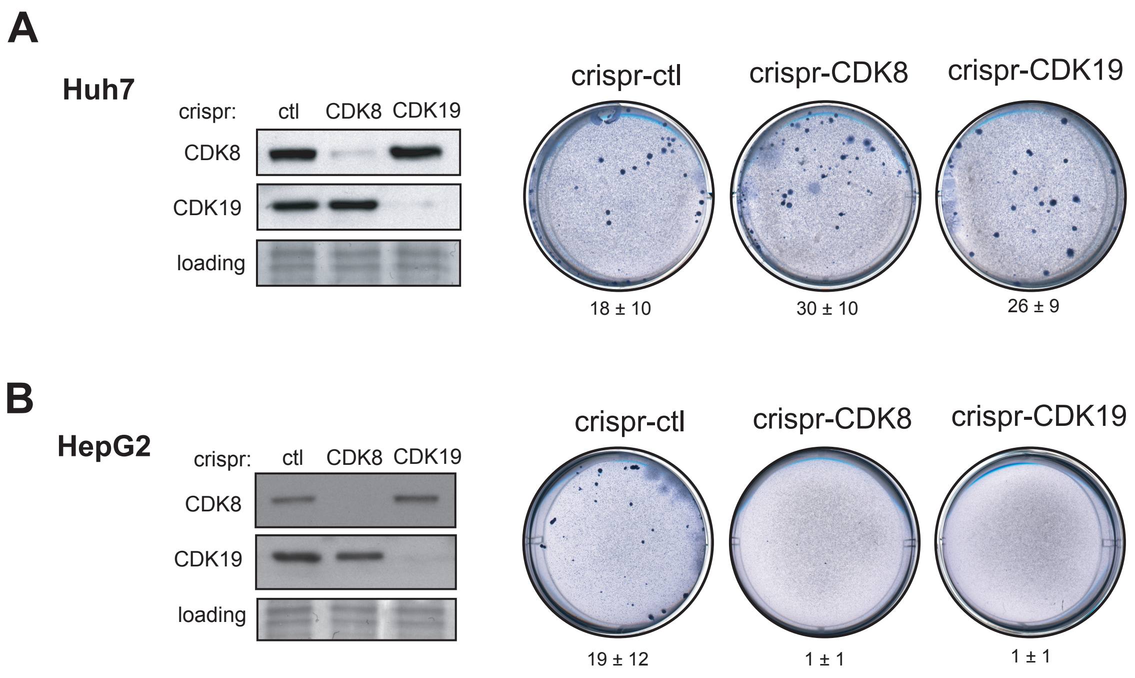


**SuppFigure 4: CDK8 and CDK19 requirement for transformation in human hepatic cancer cell lines** (**A**) Western-blot characterization of CDK8 or CDK19 removal by crispr/-cas9 gene editing and consequences on growth in soft agar for Huh7 cells. Mean ± SD number of colonies per well from three independent experiments are indicated. Note that CDK8 or CDK9 removal has no effect on growth in anchorage-independent conditions (**B**) Western-blot characterization of CDK8 or CDK19 removal by crispr/cas9 gene editing and consequences on growth in soft agar for HepG2 cells. Mean ± SD number of colonies per well from three independent experiments are indicated.
